# RobusTAD: A Tool for Robust Annotation of Topologically Associating Domain Boundaries

**DOI:** 10.1101/293175

**Authors:** Rola Dali, Guillaume Bourque, Mathieu Blanchette

## Abstract

**Motivation:** Topologically Associating Domains (TADs) are chromatin structures that can be identified by analysis of Hi-C data. Tools currently available for TAD identification are sensitive to experimental conditions such as coverage, resolution and noise level.

**Results:** Here, we present RobusTAD, a tool to score TAD boundaries in a manner that is robust to these parameters. In doing so, RobusTAD eases comparative analysis of TAD structures across multiple heterogeneous samples.

**Availability:** RobusTAD is implemented in R and released under a GPL license. RobusTAD can be downloaded from https://github.com/rdali/RobusTAD and runs on any standard desktop computer.

**Supplementary information:** Supplementary data are available at *Bioinformatics* online.

## Introduction

Topologically Associating Domains (TADs) are self-interacting genomic regions that can be identified through the analysis of data from Hi-C experiments. They have been described as key structures in chromatin organization and have been associated with roles in genome organization and gene regulation. Several tools currently exist for TAD detection, such as DomainCaller (Dixon, et al., 2012), TopDom (Shin, et al., 2016), TADbit (Serra, et al., 2017), Arrowhead (Rao, et al., 2014), HiCSeg (Lévy-Leduc, et al., 2014), Armatus (Filippova, et al., 2014), and TAD-tree (Weinreb and Raphael, 2015) to name a few. We have recently shown that the output of many of these tools is sensitive to various experimental parameters, such as sequencing coverage and binning resolution (Dali and Blanchette, 2017). Another inconvenience is often the lack of a comprehensive analysis for every bin in the genome; most tools output their significant TAD calls without providing any information about regions deemed non-significant. This leads to difficulties in reliably comparing TAD structures across different samples, for example to determine changes in TAD structures during cellular differentiation. To address these issues, we developed RobusTAD, a TAD boundary scoring tool that provides (i) TAD boundary scores for every bin in the genome and (ii) a list of most significant TAD boundaries. Our approach was created keeping several design principles in mind:

1. Robustness to coverage and noise level: TAD calls (numbers and positions) should not be too sensitive to these experimental parameters.
2. Stability at different resolution: TAD calls should remain largely unchanged if the same data is analyzed at different levels of resolution.
3. Robustness to TAD nesting and overlap: Predictions should not rely on assumptions regarding the relationships among TADs.
4. Suitability for differential TAD boundary analysis: The output of the program should allow easy comparison of TADs in Hi-C data obtained under different experimental conditions.

## Methods

A complete description of the algorithm is available in Supplementary File 1. Briefly, for each bin of an input interaction frequency matrix, the program computes left and right boundary scores. Right boundary scores are obtained for bin *b* by comparing IF values lying to the left of *b* to those spanning *b*, stratified based on pairwise distances; left boundary scores are computed symmetrically. Discrete left and right boundaries are then identified as local maxima of the boundary scores, provided they exceed a certain minimum threshold.

## Results

Fig. 1A shows RobusTAD’s boundary scores for a 3 Mb region of human chromosome 10, for GM12878 Hi-C data (Rao, et al., 2014), at different resolutions and coverage levels (obtained by down-sampling read pairs). TAD boundaries are precisely and consistently identified, in a manner that is robust to both experimental parameters. Evaluated against a set of manually annotated TADs (Dali and Blanchette, 2017), RobustTAD boundary predictions obtain positive predictive values comparable to other tools, but significantly better sensitivity (Fig. 1B). RobusTAD runs quickly on standard desktop computers; the analysis of Rao et al.’s whole genome data set at 50 kb resolution took less than 20 minutes, requiring less than 1 Gb of RAM.

**Fig. 1.**
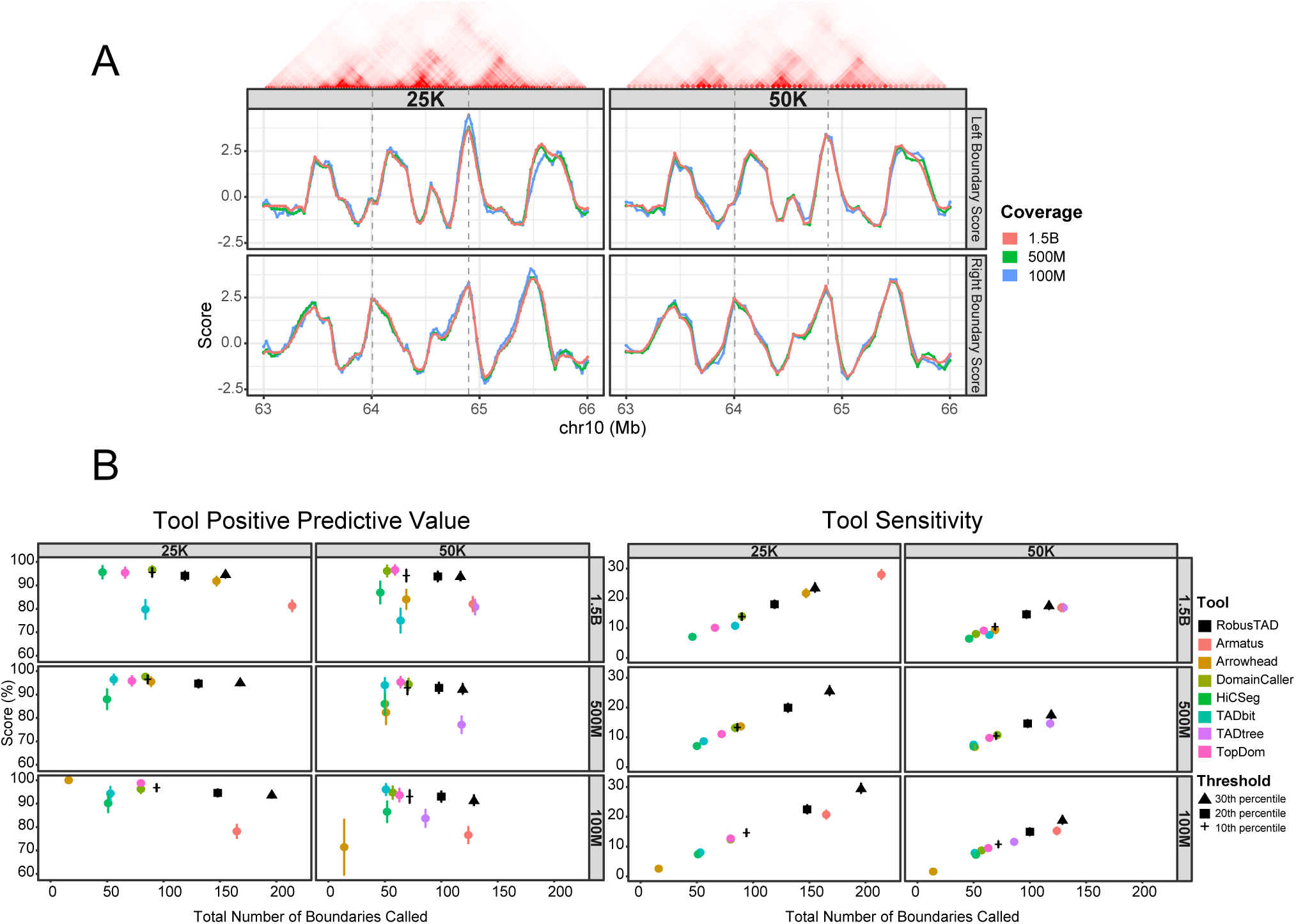
(A) Left and right TAD score profiles for chr10:63,000,000-66,000,000, at two different resolutions (25 kb and 50 kb bins) and three different levels of sequencing coverage (100, 500, and 1,500 Million read pairs). (B) Positive predictive value (left) and sensitivity (right) of RobusTAD (at three different score thresholds: top 30^th^, 20^th^ and 10^th^ percentiles) and selected TAD discovery tools against a manually annotated dataset (10 regions of 5 Mb each (Dali and Blanchette, 2017)). Error bars correspond to one std. dev. of the estimates.

## Acknowledgements

We thank Christopher Cameron for useful discussions.

## Funding

This work has been supported by a Discovery Grant to MB from the National Science and Engineering Research Council of Canada.

## Supplementary Methods

### TAD boundary scores computation

Given an interaction frequency (IF) matrix, RobusTAD first performs a simple ICE-like normalization (Imakaev, et al., 2012), if the input is not already normalized. It then computes left and right TAD boundary scores for each bin of the normalized IF matrix. We explain here how the right boundary scores are computed (see Fig. S1); the left boundary scores are calculated symmetrically. The value of RightBoundaryScore(*i*) measures the evidence for a right TAD boundary between bins *i-1* and *i*; this score will be large when a TAD ends at between bins *i-1* and *i*. This is obtained by comparing the interaction frequencies among pairs of bins (*b,b’*) lying to the left of *i* (*i-w-1 ≤ b < b’ < i*; blue triangle in Fig. 1A) to those spanning *i* (*b < i, i ≤ b’ ≤ i+w;* green triangle). Because IF values tend to decrease with genomic distance, it is important that this comparison be made keeping distances *b’-b* constant. To this end, define the IF multisets 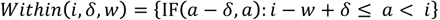, which contains the IF observed on the δ-th matrix diagonal, involving regions located up to *w* bins to the left of *i.* Define also 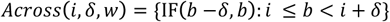, which contains interaction on the δ-th diagonal and spanning *i*. We then obtain

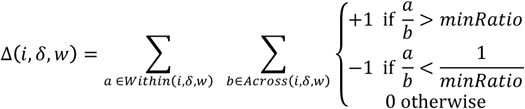

and 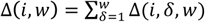, which is an integer that ranges from 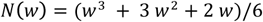 (for a perfect TAD right boundary) to −*N*(*w*). The robustness of RobusTAD owes much to the use of this count-based approach, which is quite insensitive to noise and low coverage. We normalize Δ(*i,w*) to obtain 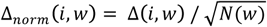. Instead of fixing *w* to a set value, we choose *w* such that Δ_*norm*_(*i,w*) is maximized, for *w* in the user-defined range *w*_*min*_,…, *w*_*max*_:

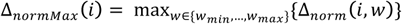. Finally, to ensure comparability across data sets, boundary scores are z-score normalized:

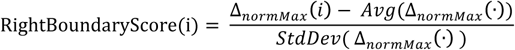

The parameters *minRatio*, *w_min_*, *and w*_max_ are user-defined. *MinRatio* (default = 1.5) controls the required contrast between a TAD and its neighbourhood: setting it to 1 allows the identification of more subtle TAD boundaries, while setting minRatio >2 selects for very sharp boundaries. *minRatio*, *w_min_*, *and w*_*max*_ control the smallest and largest genomic distance considered for the boundary score calculation. They are set by default to 250 kb and 500 kb respectively, although the results are quite insensitive to the values of those parameters. Note this does not prevent the identification of boundaries for TADs larger than *w_max_*.

### TAD boundary prediction

If the user is interested in obtaining a discrete set of sites identified as TAD boundaries, RobusTAD will call significant TAD boundaries by locating local maxima along the left and right TAD boundary score profiles that lie in the top *T* percentiles of the data (default: *T*=20%).

**Supplementary Fig. 1.**
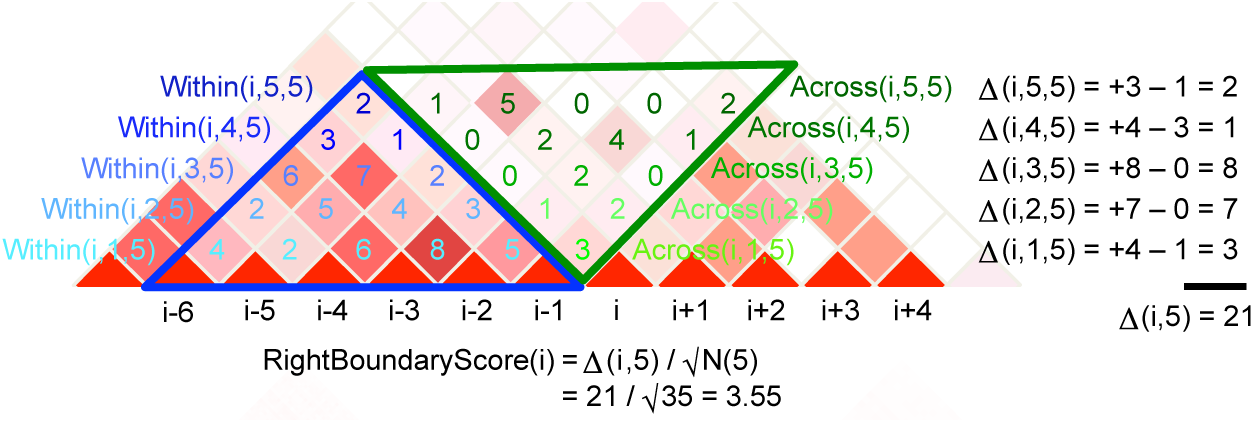
Illustration of the RightBoundaryScore calculation, for *w* = 5. Note that in reality, all values *w* ∈ {*w*_*min*_,…, *w*_*max*_} are considered, to identify the value that maximizes Δ_*norm*_(*i,w*).

